# Laboratory surrogate markers of residual HIV replication among distinct groups of individuals under antiretroviral therapy

**DOI:** 10.1101/416099

**Authors:** L.B. Giron, S.B. Tenore, L.M. Janini, M.C. Sucupira, R.S. Diaz

## Abstract

**Background:** Residual HIV-1 replication among individuals under antiretroviral therapy (ART) relates to HIV micro-inflammation.

**Objectives:** To determine levels of residual HIV replication markers among distinct subgroups of antiretroviral-treated individuals.

**Methods:** 116 patients were distributed into 5 treatment groups: first-line suppressive ART with non-nucleoside reverse-transcriptase inhibitor (NNRTI) (n=26), first-line suppressive ART with boosted protease inhibitors (PI-r) (n=25), salvage therapy using PI-r (n=27), salvage therapy with PI-r and raltegravir (n=22) and virologic failure (n=16). Episomal and total DNA quantitation was evaluated. ELISA was used for HIV antibody and LPS quantitation.

**Results:** Episomal DNA was positive in 26% to 38% of individuals under suppressive ART, being higher among individuals experiencing virologic failure (p=0.04). HIV proviral load was higher among patients with detectable episomal DNA (p=0.01). Individuals receiving initial PI-r treatment presented lower HIV antibodies (p=0.027) and LPS (p=0.029) than individuals receiving NNRTI. There was a negative correlation between episomal DNA quantitation and suppressive ART duration (p=0.04), CD4+ T-cell count (p=0.08), and CD8+ T-cell count (p=0.07).

**Conclusions:** Residual HIV replication has been inferred among individuals under suppressive ART according to episomal DNA detection. Residual replication may decrease with longer periods of suppressive ART and higher levels of CD4+ and CD8+ T cells. The relationship between episomal DNA and total DNA suggests a replenishment of the proviral reservoir with impacts on HIV persistence. Lower antibody and LPS levels among patients with initial PI-r ART suggest these regimens may more effectively suppress HIV with higher capacity to decrease the HIV antigenic component.

## Introduction

The deleterious effects of HIV are directly related to viral replication, which leads to inflammatory processes, such as the activation of CD4+ and CD8+ T lymphocytes (1). Maintaining viral replication at lower levels is critical for the reduction of cellular activation and co-morbidities related to HIV-1 infection. However, the antiretroviral therapy (ART) currently used does not completely suppress viral replication. Up to 80% of patients with undetectable viral loads according to commercial tests show an average of 3.1 copies/mL of residual viral load when ultrasensitive tests are used (2, 3). Although the stability of episomal DNA is not completely understood, extrachromosomal DNA is useful as a surrogate marker of HIV-1 replication when the HIV viral load is not detectable by currently available methods (4). Other markers that relate to HIV-1 replication among individuals under ART include proviral HIV DNA (5) and the quantitation of HIV antibody levels (6) or markers that relate to bacterial translocation (7). ART regimens differ in potency as well as in the distinct genetic barriers they create or effects they have in each step of the HIV replication cycle to alter viral dynamics. For this reason, the evaluation of circular HIV DNA could be used as a tool to indirectly compare the effectiveness of these distinct regimens on residual HIV replication. Therefore, this study aimed to analyze surrogate markers of the residual replication rates of HIV-1 among individuals receiving different antiretroviral regimens. We hypothesize that drugs from different classes and previous ART virologic failure will affect surrogate markers of HIV residual replication.

## Methods

### Patients

Patients were chosen between 2011 to 2013 in São Paulo Brazil according to their current antiretroviral treatment (see Supplementary Table 1). Individuals were under ART with undetectable plasma viral loads for at least one year, except for the virologic failure group. This study was approved by the Ethics Committee in Research at the Federal University of São Paulo (approval #0201/11), and informed consent was obtained from all patients.

**Table 1:**
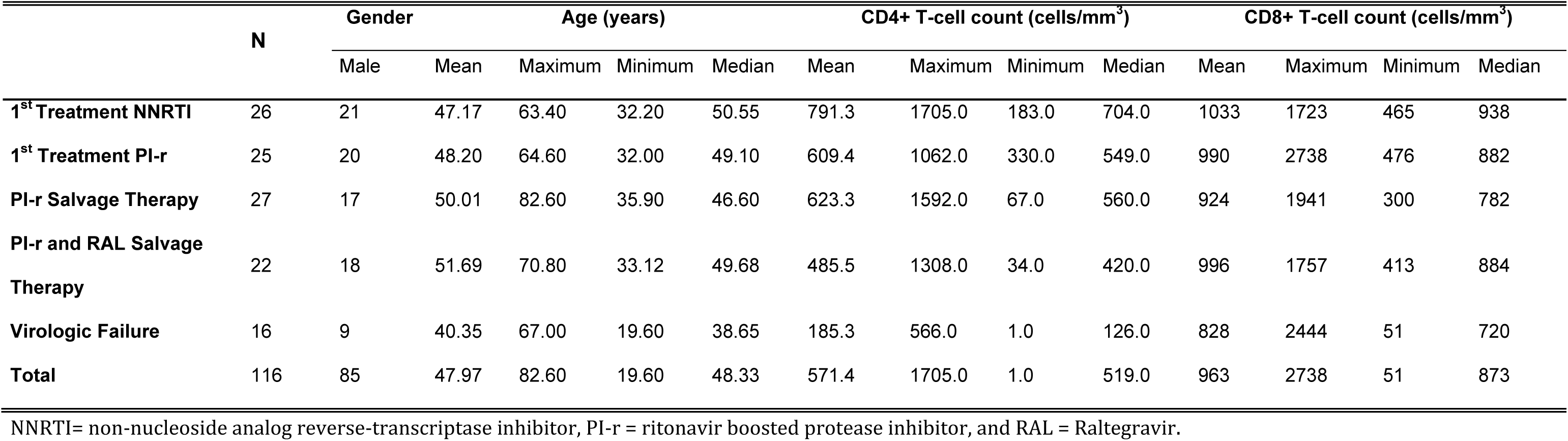
Demographic characteristics and mean T-cell counts between the different treatment groups. NNRTI= non-nucleoside analog reverse-transcriptase inhibitor, PI-r = ritonavir boosted protease inhibitor, and RAL = Raltegravir.

One hundred sixteen patients were allocated to five treatment subgroups as follows: (*1*) patients treated with two nucleoside-analogue reverse-transcriptase inhibitors (NRTI) associated with the non-nucleoside analog reverse-transcriptase inhibitors (NNRTI) Efavirenz or Nevirapine as the first ART regimen; *(2)* patients treated with two NRTIs and a protease inhibitor boosted with ritonavir (PI-r) as the first ART regimen; *(3)* patients on salvage therapy with two NRTIs and a PI-r; *(4)* patients under salvage therapy containing two NRTIs, PI-r and the integrase inhibitor raltegravir; and *(5)* patients under antiretroviral virologic failure with the confirmed presence of HIV ART resistant strains. Peripheral blood samples were collected, and clinical data on the patients were analyzed, including CD4+ and CD8+ T-cell counts, the duration of treatment with undetectable viral loads and the number of ART schemes previously used by the patient.

### HIV-1 Episomal DNA Detection and Quantitation

To obtain HIV episomal DNA, 400 μL of peripheral blood mononuclear cells (PBMC) isolated using density gradient centrifugation were extracted using a QIAprep Spin Miniprep commercial kit (Qiagen, Valencia, California, USA). After extraction, qPCR amplification was performed in a single round of 45 qPCR cycles to amplify extrachromosomal DNA as previously described (8, 9). The qPCR quantitation values were normalized based on cell numbers estimated by CCR5 quantitation and are expressed as the number of DNA copies per 10^6^ PBMC.

### Total HIV DNA Quantitation

Total viral DNA was extracted from 50 μL of PBMC using a Blood QIAamp DNA Mini Kit Mini Kit (Qiagen, Valencia, California, USA) according to the manufacturer’sinstructions. Total HIV DNA was qPCR amplified using a mix containing 1x TaqMan Universal PCR Master Mix (Applied Biosystems), 0.4 µM primers/probe(10) (F522-43 GCCTCAATAAAGCTTGCCTTGA, R626-43 GGGCGCCACTGCTAGAGA and Probe CCAGAGTCACACAACAGACGGGCACA) and 5 µl of extracted DNA. CCR5 was also used to quantify genomes to express the measurements as copies per 10^6^ PBMC.

### Quantitation of anti-HIV-1 Antibodies

HIV-1 specific antibodies were measured using the capture enzyme immunoassay kit Aware BED Incident HIV-1 EIA Test (Calypte Biomedical Corporation, Portland, Oregon, USA) according to the manufacturer’s instructions. The optical density values of the samples were normalized based on the controls (negative, calibrator, lower positive and higher positive) using the spreadsheet available at http://www.calypte.com/aware_BED.html. Specimens with an ODn > 0.8 are considered positive.

### Levels of LPS in Plasma

The quantitation of endotoxin was performed using a Limulus Amebocyte Lysate (LAL) QCL-1000 (Lonza, Walkersville, MD) kit according to the manufacturer’s instructions. The absorbance was determined spectrophotometrically at 405–410 nm. Since this absorbance is in direct proportion to the amount of endotoxin present, the concentration of endotoxin was calculated from a standard curve. The background color of the sample was subtracted.

### Statistical analysis

The Statistical Program for the Social Sciences, version 18.0 (SPSS 18.0) was used for data analysis. Descriptive analyses, ANOVA using z-score normalized data, and chi-squared tests, at a confidence level of 5%, were performed.

## Results

### Episomal DNA

The general patient data including age, gender, CD4+ and CD8+ T-lymphocyte counts, treatment time, number of regimens and number of medications used were compiled and are shown in Tables 1 and 2, grouped according to the type of ART received.

**Table 2:**
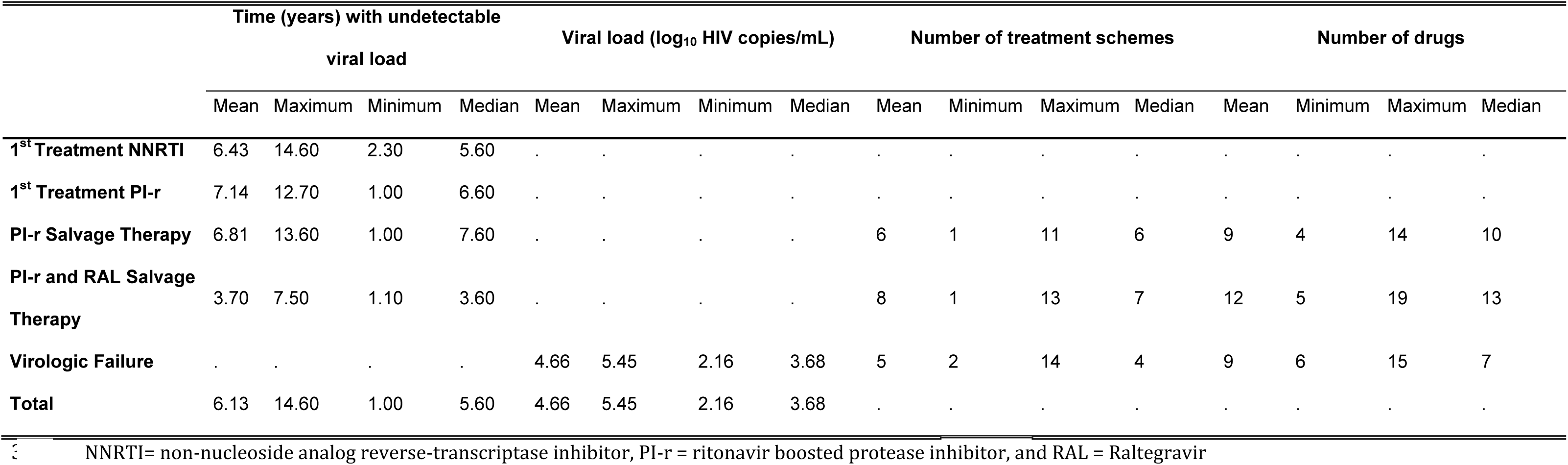
Treatment response and characteristics between the distinct treatment groups. NNRTI= non-nucleoside analog reverse-transcriptase inhibitor, PI-r = ritonavir boosted protease inhibitor, and RAL = Raltegravir

|  | Time (years) with undetectable viral load |  |  |  | Viral load (log<sub>10</sub> HIV copies/mL) |  |  |  | Number of treatment schemes |  |  |  | Number of drugs |  |  |  |
| --- | --- | --- | --- | --- | --- | --- | --- | --- | --- | --- | --- | --- | --- | --- | --- | --- |
|  | Mean | Maximum | Minimum | Median | Mean | Maximum | Minimum | Median | Mean | Minimum | Maximum | Median | Mean | Minimum | Maximum | Median |
| **1<sup>st</sup> Treatment NNRTI** | 6.43 | 14.60 | 2.30 | 5.60 | . | . | . | . | . | . | . | . | . | . | . | . |
| **1<sup>st</sup> Treatment PI-r** | 7.14 | 12.70 | 1.00 | 6.60 | . | . | . | . | . | . | . | . | . | . | . | . |
| **PI-r Salvage Therapy** | 6.81 | 13.60 | 1.00 | 7.60 | . | . | . | . | 6 | 1 | 11 | 6 | 9 | 4 | 14 | 10 |
| **PI-r and RAL Salvage Therapy** | 3.70 | 7.50 | 1.10 | 3.60 | . | . | . | . | 8 | 1 | 13 | 7 | 12 | 5 | 19 | 13 |
| **Virologic Failure** | . | . | . | . | 4.66 | 5.45 | 2.16 | 3.68 | 5 | 2 | 14 | 4 | 9 | 6 | 15 | 7 |
| **Total** | 6.13 | 14.60 | 1.00 | 5.60 | 4.66 | 5.45 | 2.16 | 3.68 | . | . | . | . | . | . | . | . |

2-LTR circles were detected in 39 (34%) of the patients in the study. Table 3 summarizes the measurements obtained according to treatment group. The treatment group had no effect on the quantitation of episomal HIV DNA (F (115,4) = 1.263, p = 0.289). The prevalence of detectable 2-LTR (n=39) was not different between the groups (F(38,4)=1.014, p=0.414).

**Table 3:**
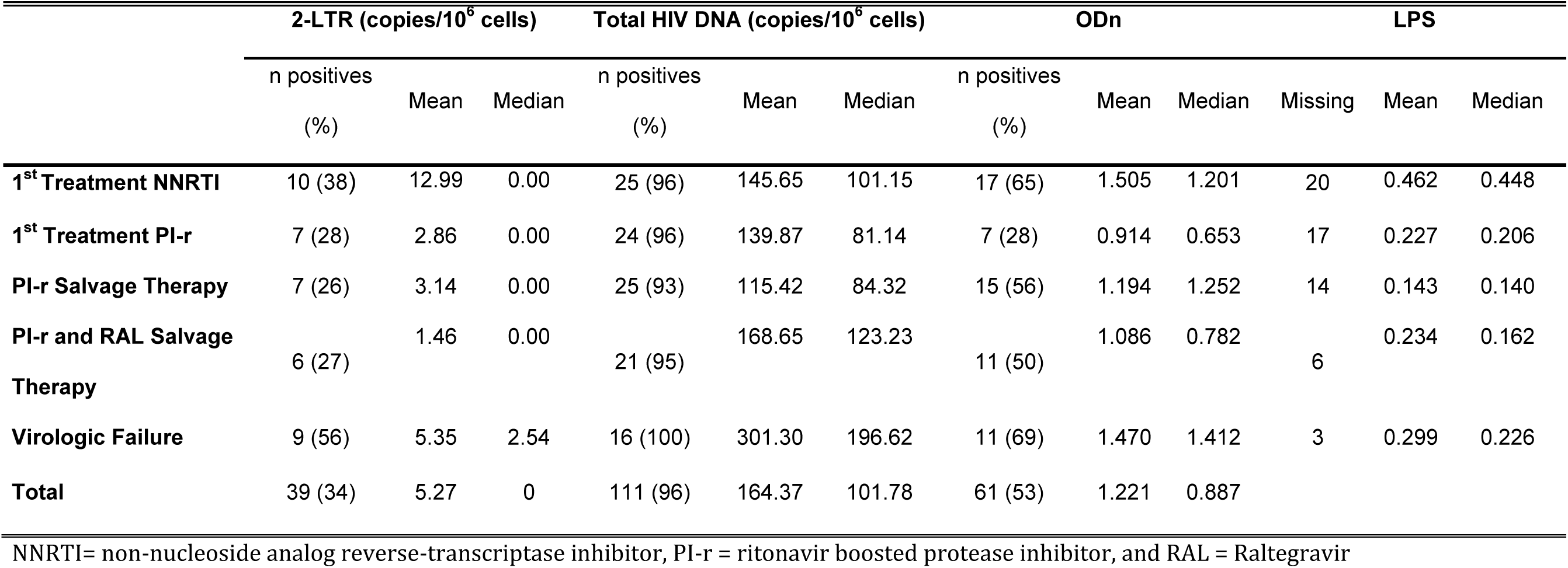
Prevalence of 2-LTR and HIV total DNA and EIA Optical density (ODn) and plasma LPS for the different treatment groups NNRTI= non-nucleoside analog reverse-transcriptase inhibitor, PI-r = ritonavir boosted protease inhibitor, and RAL = Raltegravir

|  | 2-LTR (copies/10<sup>6</sup> cells) |  |  | Total HIV DNA (copies/10<sup>6</sup> cells) |  |  | ODn |  |  | LPS |  |  |
| --- | --- | --- | --- | --- | --- | --- | --- | --- | --- | --- | --- | --- |
|  | n positives (%) | Mean | Median | n positives (%) | Mean | Median | n positives (%) | Mean | Median | Missing | Mean | Median |
| **1<sup>st</sup> Treatment NNRTI** | 10 (38) | 12.99 | 0.00 | 25 (96) | 145.65 | 101.15 | 17 (65) | 1.505 | 1.201 | 20 | 0.462 | 0.448 |
| **1<sup>st</sup> Treatment PI-r** | 7 (28) | 2.86 | 0.00 | 24 (96) | 139.87 | 81.14 | 7 (28) | 0.914 | 0.653 | 17 | 0.227 | 0.206 |
| **PI-r Salvage Therapy** | 7 (26) | 3.14 | 0.00 | 25 (93) | 115.42 | 84.32 | 15 (56) | 1.194 | 1.252 | 14 | 0.143 | 0.140 |
| **PI-r and RAL Salvage Therapy** | 6 (27) | 1.46 | 0.00 | 21 (95) | 168.65 | 123.23 | 11 (50) | 1.086 | 0.782 | 6 | 0.234 | 0.162 |
| **Virologic Failure** | 9 (56) | 5.35 | 2.54 | 16 (100) | 301.30 | 196.62 | 11 (69) | 1.470 | 1.412 | 3 | 0.299 | 0.226 |
| **Total** | 39 (34) | 5.27 | 0 | 111 (96) | 164.37 | 101.78 | 61 (53) | 1.221 | 0.887 |  |  |  |

There was no difference in the quantitation of 2-LTR circles among groups with first treatment (F(49,1) = 1.429, p = 0.23, Figure 1A). Additionally, there was no difference (F (47,1) = 1.692, p = 0.20) when comparing the 2 distinct salvage therapy groups. We also observed no difference between the two groups receiving PI-r (F(50,1)=0.197, p=0.65). Furthermore, there was no significant difference between the first treatment groups together and the salvage groups together, (F (98,1) = 1.229, p = 0.27) nor when comparing the groups with virologic success to that of virologic failure (F (114,1) = 0.601, p = 0.44).

**Figure 1:**
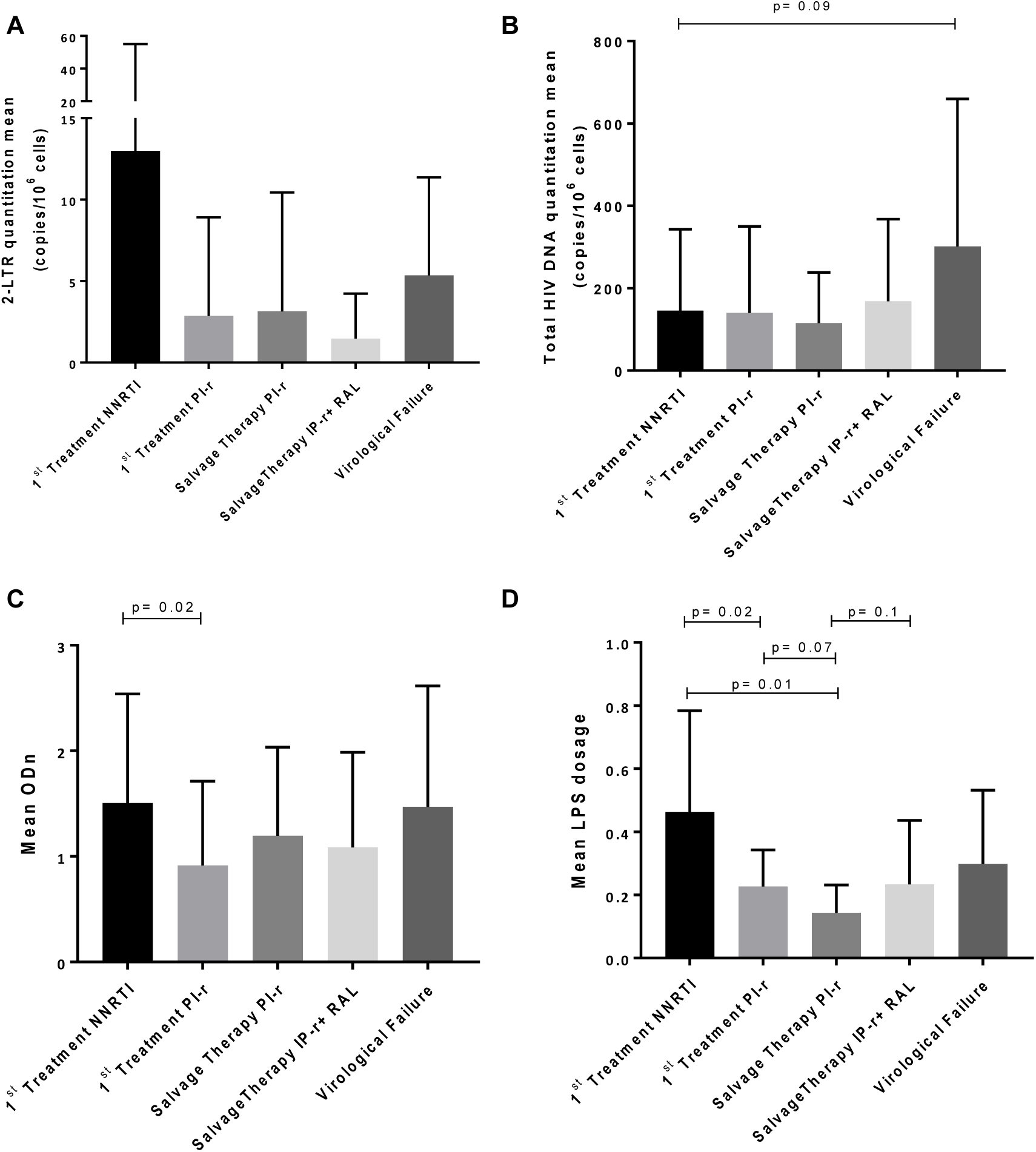
Mean levels of 2-LTR HIV DNA (Panel A), total HIV DNA (Panel B), EIA optical density (ODn; Panel C) and LPS levels (panel D) among the different treatment groups. Bars show standard deviation. P values lower than 0.1 are indicated.

We then transformed episomal DNA quantitation into a categorical variable for detection and named samples LTR positive when detection was possible and LTR negative when there was no detection. Based on this categorization, we performed a chi-square test. The results showed no statistically significant association between the received treatment and the detection of circular DNA (χ^2^ (3) = 5.412, p = 0.248). Comparing the number of positive episomal DNA samples between the subjects with virologic suppression and individuals experiencing virologic failure, there was an increase in the number of episomal DNA-positive samples in the failure group (χ^2^ (3) = 4.259, p = 0.039, Figure 2A). In addition, the mean of total DNA was higher among individuals with positive episomal DNA (ANOVA, F (109,1) = 2.794, p = 0.09; Figure 2B).

**Figure 2:**
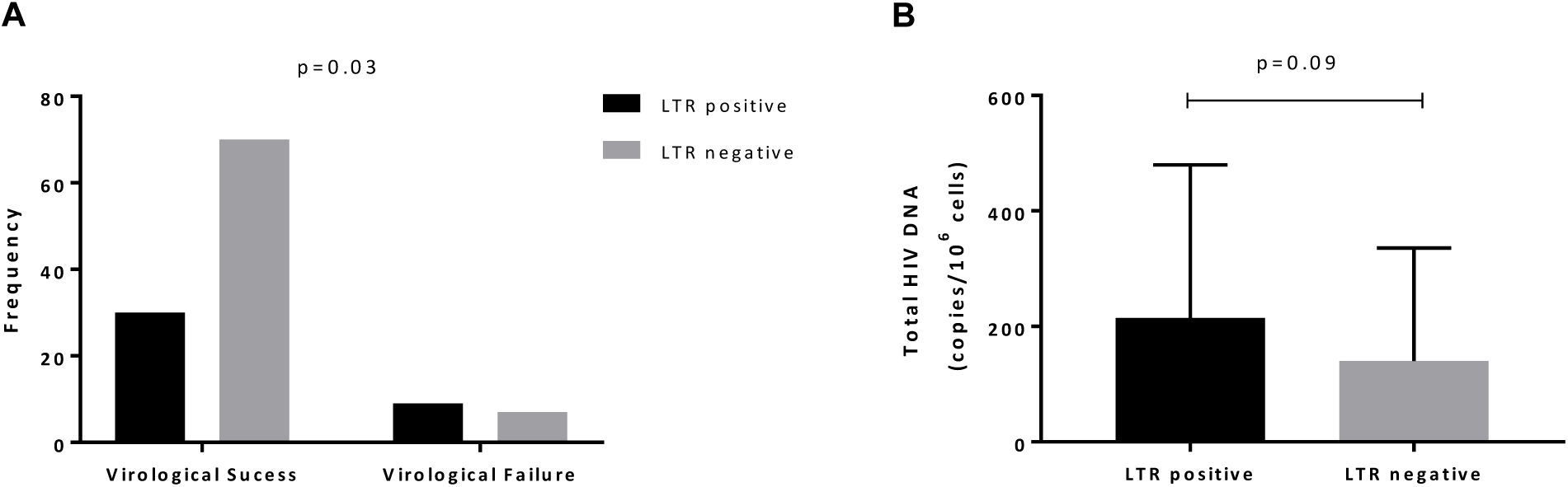
(A) Frequency of 2-LTR positive samples among patients with virologic success and virologic failure. (B) Mean levels of total HIV DNA in 2-LTR positive and negative samples. Bars show standard deviation.

### Total HIV DNA

Total HIV DNA was detected in 111 (96%) of the patients included in the study (Table 3). An ANOVA tests showed no differences between the treatment groups for quantitation of total HIV DNA (F (115,4) = 2.015, p = 0.098; Table 3 and Figure 1B). Additionally, there was no difference in total HIV DNA quantitation between the groups with a first-line regimen (Groups 1 and 2, F(47,1) = 0.010, p = 0.922), nor between the two groups on salvage therapy (Groups 3 and 4, F (44,1) = 1.230, p = 0.273), nor between the groups on a first-line regimen and salvage therapy (F (93,1) = 0.007, p = 0.935). Finally, there was a difference between the groups with virologic success and virologic failure (F (109,1) = 7.528, p = 0.007) in which virological failure group shows higher total HIV DNA mean.

There was no statistical significance between total HIV DNA and the other tested variables.

### Quantitation of anti-HIV-1 antibodies

In this test, we considered samples with normalized optical densities (ODn) higher or equal to 0.8 as positive. Table 3 summarizes the measurements obtained according to treatment group. ANOVA showed no differences in the HIV antibody levels between the groups (F (115,4) = 1.675, p = 0.161, Figure 1C). However, the antibody levels were higher among patients given first treatment with NNRTI compared to first treatment with PI-r (ANOVA; F (49,1) = 5.189, p = 0.027). There was no difference when comparing the two types of salvage therapy schemes (F(47,1)=0.189, p = 0.66) nor between the first-line treatment and salvage therapy groups (F(98,1)=0.146, p = 0.70). In addition, there was no difference when comparing the groups with virologic successful and virologic failure (F(114,1)=1.289, p = 0.25).

Considering ODn as a categorical variable in which positive samples had an ODn ≥ 0.8, there was a decreased number of positive samples in the first treatment group using PI-r (χ ^2^ (1) = 9.600, p = 0.007) compared to the first treatment group using NNRTI as well as an increase in positivity when compared to salvage therapy with PI (χ ^2^ (1) = 4.038, p = 0.044) (Figure 3). There was no significant difference between the first-line regimen groups and the salvage therapy groups, (χ ^2^ (1)=0.360, p=0.34) nor any difference when comparing groups with or without virologic failure (χ ^2^ (1)=1.945, p=0.13).

**Figure 3:**
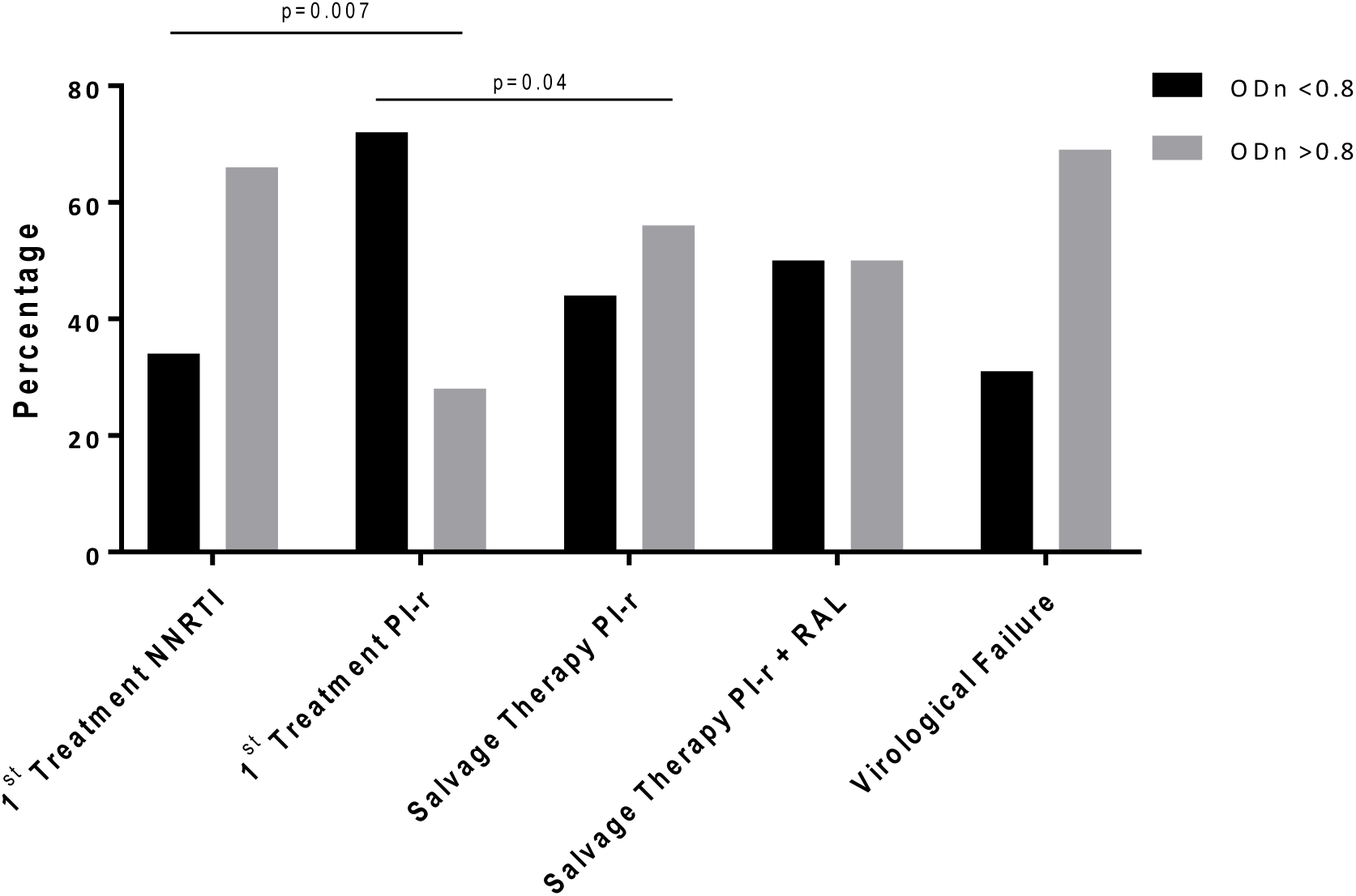
Frequency of samples in which the less sensitive anti-HIV EIA optical density (ODn) was higher or lower than 0.8, indicating a positive and negative result, respectively

Positive antibody quantitation was not associated with the positivity of episomal DNA (χ ^2^ (1) = 1.889, p = 0.119) or with the episomal DNA quantitation (F(114,1)=0.112, P=0.738). Patients with positive antibody quantitation showed slightly higher HIV total DNA (F(109,1)=2.787, p=0.09).

### LPS quantitation

Due to the unavailability of samples, LPS quantitation was performed for only 55 patients (Table 3). An ANOVA test showed a significant difference between the first-line regimen groups, with LPS higher among individuals treated with NNRTI compared to PI-r (F(55,4)=2.947, p= 0.029, Figure 1D), as well as between the NNRTI and salvage therapy groups and the PI-r group (p=0.019, Bonferroni Test).

### Correlations

Spearman correlation tests were performed only with samples in which episomal DNA was detected. There was a negative correlation between the quantitation of episomal DNA and the CD8+ T-cell count (ρ = −0.426, p = 0.007) and the CD4+ T-cell count (ρ = - 0.276, p = 0.08), LPS quantitation in plasma (ρ = −0.500, p = 0.041) and treatment time with an undetectable viral load (ρ = −0.358, p = 0.044).

Spearman correlation between total HIV DNA showed positive correlation with episomal DNA quantitation (ρ = 0.256, p = 0.007), antibody levels (ρ = 0.181, p = 0.05) and also a negative correlation with CD8+ T-cell count (ρ = −0.243, p = 0.01) and trend to correlate with CD4+ T-cell count (ρ = −0.16, p = 0.09).

## Discussion

As mentioned before, antiretroviral treatment is not fully suppressive in all individuals, as shown by the detection of viremia in individuals evaluated with ultrasensitive viral load assays (3) or with tests for cell-associated RNA (11). Interestingly, this residual viremia may come from so-called sanctuaries, such as the gastrointestinal tract (12). As such, they form an obstacle for achieving a sterilizing cure. Furthermore, specific HIV inflammation inferred by the levels of T-cell lymphocyte activation persists among antiretroviral treated individuals in spite of undetected viral loads (13). Efforts and strategies to mitigate HIV-related inflammation is currently a major task. One effective way to decrease this inflammation would be to maximize the antiretroviral suppressive effect, thus reducing residual replication.

Furthermore, continuous suppressive therapy is able to decrease the number of latent HIV infected cells over time (14), bringing the individual close to a sterilizing cure when the right strategies become available. On the other hand, residual viremia is conceivably able to replenish latent HIV reservoirs.

To learn more about residual HIV-1 replication among individuals under ART, we used different surrogate markers of HIV replication. The presence and quantitation of episomal HIV DNA has been considered one accurate marker to infer active HIV replication and its entrance into the cell environment (9, 15). Total or integrated HIV DNA also indicates the size the HIV infected cell pool. It is well known that early treatment initiation affects the number of latently infected cells (16), and over time, cells will exit latency and die, decreasing the proviral DNA pool in ART treated individuals. The levels of HIV-1 antibodies detected using less sensitive assays also relate to the levels of HIV-1 replication (6). As HIV-1 residual replication may come from the gastrointestinal tract (12), it is also conceivable that less effective antiretroviral treatment could be associated with higher levels of bacterial translocation (7) and therefore increasing laboratory translocation markers such as LPS or sCD14 levels.

We also wanted to investigate the relationship between different HIV ART schemes or strategies. The main questions were: is initial treatment more suppressive when two distinct steps of reverse transcription are inhibited, such as schemes using NRTIs with an NNRTI, or is the inhibition of pre- and post-integration more effective, such as schemes using NRTIs and boosted PI? A number of clinical trials comparing NNRTI with boosted PIs as the second antiretroviral class show one advantage of NNRT, which relies mainly on tolerance and adherence issues, since boosted PI schemes do not present antiretroviral resistance upon failure (17). The other question is whether salvage therapy is associated with more residual HIV replication than initial antiretroviral therapy. Usually, salvage therapy relies on a boosted PI-based regimen with or without the use of a new antiretroviral class. Therefore, a further question would be whether the association of a third antiretroviral class would more suppressive than salvage therapy schemes containing 2 NRTIs and a boosted PI only. We therefore performed a cross-sectional evaluation of a distinct group of individuals under “suppressive” antiretroviral treatment with good treatment adherence using 2 NRTIs and either efavirenz/nevirapine or PI-r as the first-line treatment. We also evaluated individuals who previously experienced antiretroviral virologic failure and had their HIV viremia subsequently suppressed with 2 NRTIs and a PI-r only or PI-r associated to raltegravir. We also used as a “control group”, individuals experiencing virologic failure in which antiretroviral resistance had been detected. We attempted to avoid individuals not using or adhering to ART at the time of the study.

We were able to confirm the relationship between HIV-1 replication and the detection of episomal DNA, which was higher among individuals experiencing virologic failure compared to individuals with viral loads below detection, even with the smaller sample size of the virologic failure group.

We also detected a negative correlation between episomal DNA quantitation and the time of treatment with undetectable viral loads as well as a negative correlation between episomal DNA and CD8+ T-cell counts. It is conceivable that lower CD8 levels enable HIV-1 viral replication, as has been seen in animal models; the elimination of CD8+ T cells using monoclonal antibodies was associated with the return of detectable viremia in SIV-infected monkeys in spite of the use of suppressive ART (18). Likewise, we hypothesize that longer durations of effective antiretroviral treatment will progressively strengthen the immune system, by increasing the number of naïve CD4+ T cells and thus further decreasing residual HIV-1 replication. This speculation is further supported by the observation of a negative correlation between the levels of episomal DNA and CD4+ T-cell counts. However, we were not able to explain the negative correlation between episomal DNA levels and LPS levels.

Interestingly, the levels of total HIV DNA were found to be higher among individuals with evidence of residual HIV replication as inferred by the presence of episomal DNA. This association suggests that the pool of infected cells is being replenished or maintained in association with residual HIV replication.

We were not able to detect any differences between episomal or total DNA levels between first-line regimens and successful salvage therapy regimens, nor between NNRTI versus PI-r regimens or salvage therapy using two or three classes (NRTI + PI-r versus NRTI + PI-r and raltegravir). However, the levels of antibodies were lower in first-line PI-r ART compared to the NNRTI group as the number of negative antibody results were higher among the initial PI-r treatment group. Furthermore, the levels of LPS were higher among the NNRTI first-line treatment group compared to the first-line PI-r or other salvage therapy groups that also have a PI-r in the treatment scheme. Notably, the proportion of patients taking tenofovir, abacavir or zidovudine was similar in the PI-r and NNRTI groups (Table S1). Although clinical trials have noted that NNRTI-based regimens are usually more durable and effective than PI-r-based regimens despite a basal viral load and higher CD4+ T-cell levels, these results are mainly due to better performance of intention to treat analyses, which are influenced by tolerance and adherence issues. Importantly, 14 individuals in the PI-r group were treated with boosted atazanavir, whereas 11 were treated with boosted lopinavir [Table S1]. However, this study analyzed patients on stable ART without adherence or tolerability issues. We can therefore hypothesize that the effective inhibition of two different steps of the HIV replicative cycle is more effective than inhibiting only one step.

We recognize that the retrospective cross-sectional nature of this study may preclude more definite conclusions. The evaluation of only one time point in this group prevents us from understanding the dynamics of these surrogate markers for HIV replication. Furthermore, other sensitive assays measuring residual HIV replication, such as cell-associated RNA or inflammatory markers, have not been evaluated here.

However, we were able to clearly demonstrate that episomal DNA was present in 26% to 38% of individuals with “successful” antiretroviral treatment, thus suggesting that residual HIV replication is occurring despite the scheme analyzed here. We were also able to demonstrate the association of PI-r schemes with lower antibody and LPS levels, which deserves further confirmation to better understand the related mechanisms involved that can explain these findings.

## Acknowledgements

The authors are grateful to the study participants for their generous contribution to this research. We thank Rose Gabriel for helping with patient selection and recruitment.

